# Treatment-shortening effect of a novel regimen combining high-dose rifapentine and clofazimine in pathologically distinct mouse models of tuberculosis

**DOI:** 10.1101/556654

**Authors:** Vikram Saini, Nicole C. Ammerman, Yong Seok Chang, Rokeya Tasneen, Richard E. Chaisson, Sanjay Jain, Eric Nuermberger, Jacques H. Grosset

**Affiliations:** Center for Tuberculosis Research, Johns Hopkins University School of Medicine, Baltimore, Maryland, USA; MedStar Health Internal Medicine, Baltimore, Maryland, USA

## Abstract

High-dose rifapentine and clofazimine have each separately been associated with treatment-shortening activity when incorporated into tuberculosis (TB) treatment regimens. We hypothesized that both modifications, *i.e.,* the addition of clofazimine and the replacement of rifampin with high-dose rifapentine, in the first-line regimen for drug-susceptible TB would significantly shorten the duration of treatment necessary for cure. We tested this hypothesis in a well-established BALB/c mouse model of TB chemotherapy and also in a C3HeB/FeJ mouse model in which mice can develop caseous necrotic lesions, an environment where rifapentine and clofazimine may individually be less effective. In both mouse models, replacing rifampin with high-dose rifapentine and adding clofazimine in the first-line regimen resulted in greater bactericidal and sterilizing activity than either modification alone, suggesting that a rifapentine- and clofazimine-containing regimen may have the potential to significantly shorten the treatment duration for drug-susceptible TB. These data provide preclinical evidence supporting the evaluation of regimens combining high-dose rifapentine and clofazimine in clinical trials.

## INTRODUCTION

In its most recent global tuberculosis (TB) report, the World Health Organization (WHO) estimated that 10 million incident cases of TB occurred in 2017, highlighting that the global TB epidemic is tragically far from controlled (1). This sustained burden of TB occurs despite the existence of a highly efficacious regimen for treatment of drug-susceptible TB, which accounted for about 9.4 million or 94% of the estimated TB cases in 2017. Many factors, ranging from individual to programmatic levels, and varying widely in nature and by environment, contribute to barriers that prevent patients with TB from receiving or completing curative treatment (2); one such factor that globally impacts TB control is the duration of treatment.

The first-line regimen for treatment of drug-susceptible TB consists of daily administration of four drugs (rifampin, isoniazid, pyrazinamide, and ethambutol) for at least two months, followed by an additional four months of daily rifampin and isoniazid (3, 4). In addition to the burden on individual patients, administration of a six-month, multidrug regimen requires significant resources from a public health perspective, especially if treatment is directly observed as is recommended (5, 6). Incomplete administration can lead to treatment failure and the selection and spread of drug-resistant *Mycobacterium tuberculosis*; thus, curative regimens of shorter duration could significantly improve global TB control efforts.

In a well-established BALB/c mouse model of TB chemotherapy (7), increasing the rifamycin exposure, especially by replacing rifampin with daily high-dose rifapentine, significantly decreased the duration of treatment necessary to achieve relapse-free cure (8, 9). These relapse-based preclinical studies were among critical data supporting the evaluation of high-dose rifapentine in a phase 2 clinical trial for treatment of drug-susceptible TB (ClinicalTrials.gov identifier NCT00694629), where higher rifapentine exposures were strongly associated with higher bactericidal activity. A phase 3 trial is currently underway to determine whether replacing rifampin with high-dose rifapentine can shorten the treatment of drug-susceptible TB to four months (ClinicalTrials.gov identifier NCT02410772).

The anti-leprosy drug clofazimine has been indirectly associated with treatment-shortening when incorporated into regimens for multidrug-resistant-(MDR-) TB (10–15). In a BALB/c mouse model of MDR-TB chemotherapy, clofazimine specifically and directly contributed potent bactericidal and treatment-shortening activity to a regimen of second-line drugs (16), and in 2016, the WHO included clofazimine in its “Shorter MDR-TB Regimen,” referring specifically to the sterilizing (*i.e.*, treatment-shortening) activity that this drug may add to the regimen (17). The addition of clofazimine to the first-line regimen was also evaluated in BALB/c mice, where again it was shown to directly contribute significant activity, shortening the duration of treatment necessary to achieve relapse-free cure by up to 2 months (18, 19). The addition of clofazimine to the first-line regimen is currently being evaluated in the phase 2/3 TRUNCATE-TB study (ClinicalTrials.gov identifier NCT03474198).

Because replacement of rifampin with high-dose rifapentine in the first-line regimen and the addition of clofazimine separately added substantial treatment-shortening activity in the BALB/c mouse model, we hypothesized that combining these modifications would contribute greater treatment-shortening activity than either modification alone. To test this hypothesis, we used the BALB/c mouse model to directly compare the bactericidal and sterilizing activity of four daily regimens: (i) the first-line regimen; and the same regimen with either (ii) addition of clofazimine; (iii) replacement of rifampin with high-dose rifapentine; or (iv) the same regimen with both addition of clofazimine and replacement of rifampin with high-dose rifapentine.

The BALB/c mouse model is commonly used for preclinical studies to evaluate the efficacy of novel drug regimens (7). However, whereas human TB disease is associated with both intracellular bacteria and a large extracellular bacterial population in necrotic caseous lesions and cavities (20), the BALB/c model creates only cellular lung lesions with intracellular (mostly intra-macrophage) bacilli and specifically lacks the caseous hallmarks of human TB, raising concerns that it may not fully represent the activity of drugs and regimens across the full spectrum of human disease (7, 21–24). Both rifapentine and clofazimine are known to accumulate within macrophages (25, 26), and it has also been reported that both of these drugs do not diffuse well into caseous lesions (27–30). Therefore, it is possible that the BALB/c mouse model could overestimate the activity of these two particular drugs. To address this issue, we also evaluated the same four regimens in a C3HeB/FeJ mouse model of TB. In this model, mice can develop necrotic caseous lung lesions with a significant extracellular bacterial population and even lung cavities (7, 22–24, 31–35). The standard TB treatment regimen performs similarly in this model as in the BALB/c model (9, 22, 36–38), thus allowing for the specific evaluation of clofazimine and rifapentine activity in this disease setting.

## RESULTS

The scheme of the study is presented in Table 1. The primary outcome of this study was the proportion of mice with relapse-free cure, *i.e.,* culture-negative lungs six months after stopping treatment. The secondary endpoint was bactericidal activity, *i.e.*, the decline of *M. tuberculosis* colony-forming unit (CFU) counts during treatment.

**Table 1.**
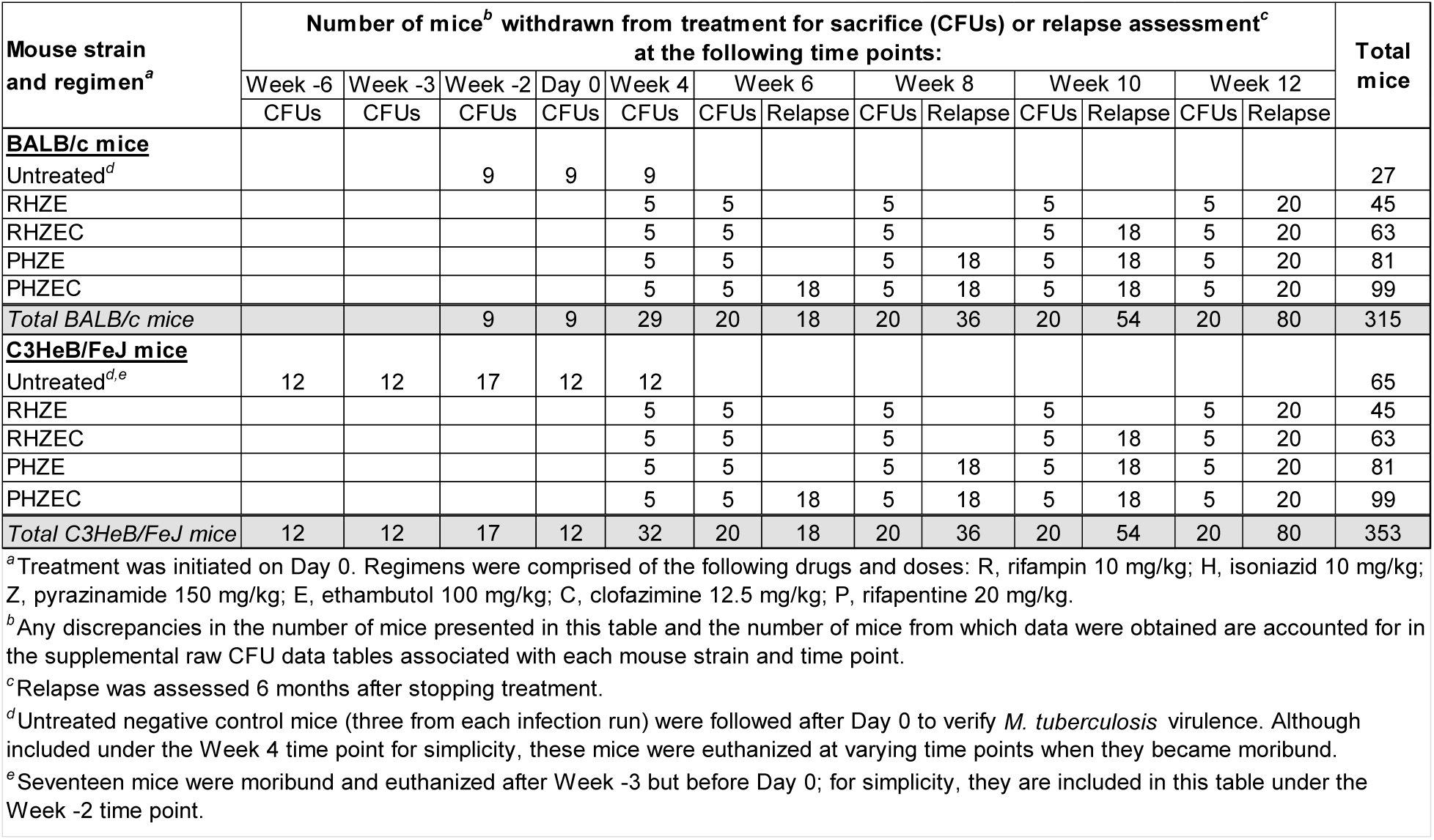
Final experiment scheme for BALB/c and C3HeB/FeJ mouse models.

### Establishment of infection: BALB/c mice

The day after aerosol infection with *M. tuberculosis* strain H37Rv, the mean bacterial implantation was 4.25 (SD 0.15) log_10_ CFU/lung (Figs. 1, **S1**; **Tables S1**, **S2**). Two weeks later, at treatment initiation, the mean bacterial burden in the lungs increased to 7.13 (SD 0.11) log_10_ CFU/lung (**Table S3**). As expected in this model (7), the untreated negative control mice became moribund three weeks after infection and were euthanized; these mice had a mean bacterial burden of 8.39 (SD 0.16) log_10_ CFU/lung (**Table S3**), and evenly distributed, small, uniform lung lesions were observed (**Fig. S2**).

**Figure 1.**
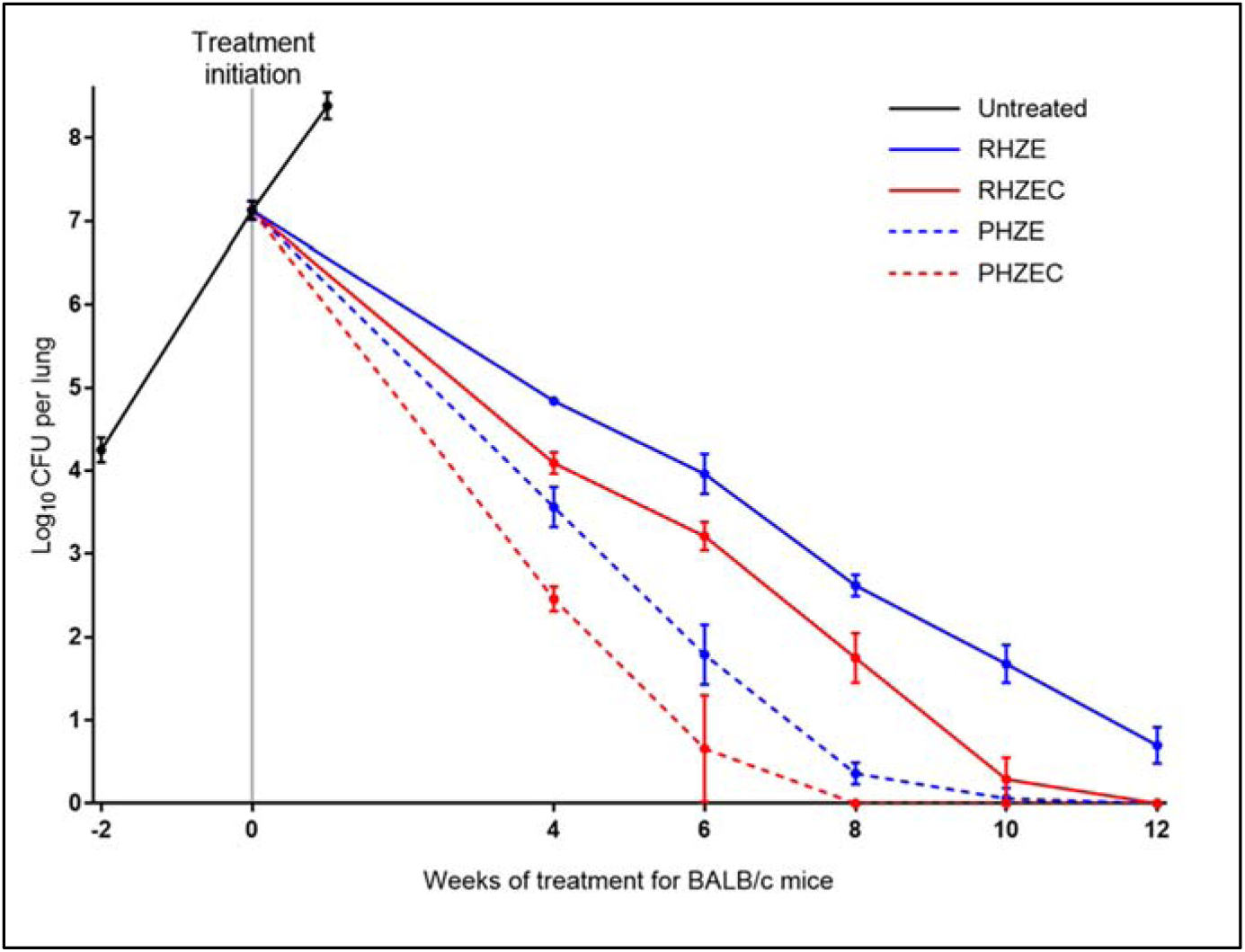
Lung CFU counts of BALB/c mice before and during treatment. Data points represent mean values, and error bars represent the SD (3-9 mice per group per time point). Treatment regimens are described in Table 1. Raw CFU data are presented in **Table S2** (Week −2/day after infection), **Table S3** (Day 0 and Untreated group), and **Tables S4**, **S5**, **S6**, **S7**, and **S8** (Week 4, 6, 8, 10, and 12, respectively).

### Assessment of bactericidal activity: BALB/c mice

The RHZE control regimen performed as expected in this model, resulting in killing of 2.3 log_10_ CFU/lung by 4 weeks, with additional killing thereafter of approximately 1 log_10_ CFU per two weeks (Fig. 1; **Tables S4-S8**). After 12 weeks of treatment, all RHZE-treated mice remained culture positive with a mean bacterial burden of 0.70 (0.22) log_10_ CFU/lung, and lung lesions were still visible (**Figs. S3-S7**). Compared to RHZE, the clofazimine-containing RHZEC regimen significantly increased the bactericidal activity by 0.8-1.0 log_10_ CFU/lung at each time point (p < 0.0001 at each time point), and all mice were culture-negative by 12 weeks. Replacement of rifampin with high-dose rifapentine had an even larger effect on the activity of the standard regimen. Administration of PHZE increased the bactericidal activity by 1.3 log_10_ CFU/lung at 4 weeks and by about 2.5 log_10_ CFU/lung at both 6 and 8 weeks compared to administration of RHZE (p < 0.0001 at each time point). Except for one mouse with a single CFU, the mice receiving PHZE were culture-negative by 10 weeks, and this regimen was also associated with significantly lower lung CFU counts than the RHZEC regimen (p < 0.01 at all time points up to 10 weeks). Despite that the lungs of PHZE-treated mice had significantly lower CFUs than the lungs of RHZEC-treated mice, gross lung lesions were consistently more visible in the PHZE-treated mice (**Figs. S3-S7**). Finally, replacement of rifampin with high-dose rifapentine plus clofazimine had the most potent bactericidal activity of all regimens tested. PHZEC was significantly more bactericidal than the PHZE regimen and all time points up to 8 weeks (p < 0.0001 at each time point), when mice receiving PHZEC became culture-negative.

### Assessment of sterilizing activity: BALB/c mice

The raw lung CFU data at relapse assessment in mice treated for 6, 8, 10, and 12 weeks are presented in **Tables S9**, **S10**, **S11**, and **S12**, respectively. As expected, nearly all mice (19/20) that received RHZE for 12 weeks were culture-positive 6 months after stopping treatment (Table 2; **Fig. S8**). Among mice treated with RHZEC for 10 weeks, nearly one-half experienced relapse, but no relapses occurred in mice that received this regimen for 12 weeks (p < 0.0001). Thirty percent of mice that received PHZE for 8 weeks relapsed, and relapse occurred in only one mouse that received this regimen for 10 weeks. Similarly, thirty percent relapse was observed in mice that received PHZEC for 6 weeks, and no relapses occurred in mice that received this regimen for 8 weeks (p = 0.09). However, there were two relapses among the mice that received PHZEC for 10 weeks, with one mouse yielding only a single CFU at relapse assessment (**Table S11**). Finally, there was no relapse after treatment for 12 weeks. Overall, the rank order of the regimens’ bactericidal activity corresponded to their sterilizing activity, *i.e.*, PHZEC > PHZE > RHZEC > RHZE. Among culture-positive mice, the bacterial burden at relapse assessment was lowest for mice that received PHZEC (all less than 4.5 log_10_ CFU/lung for all mice regardless of treatment duration), and no mouse from any treatment group relapsed with a burden greater than 5.3 log_10_ CFU/lung (**Fig. S8**).

**Table 2.** Relapse results determined 6 months after stopping treatment.

| Regimen<sup>a</sup> | Mouse strain | Proportion of mice with culture-positive relapse by treatment duration |  |  |  |
| --- | --- | --- | --- | --- | --- |
|  |  | 6 weeks | 8 weeks | 10 weeks | 12 weeks |
| RHZE | BALB/c | --- | --- | --- | 19/20 |
|  | C3HeB/FeJ | --- | --- | --- | 11/19 |
| RHZEC | BALB/c | --- | --- | 10/18 | 0/20 |
|  | C3HeB/FeJ | --- | --- | 7/17 | 3/18 |
| PHZE | BALB/c | --- | 6/20 | 1/18 | 0/18 |
|  | C3HeB/FeJ | --- | 10/18 | 0/17 | 2/20 |
| PHZEC | BALB/c | 5/18 | 0/18 | 2/18 | 0/20 |
|  | C3HeB/FeJ | 5/17 | 1/18 | 1/18 | 0/18 |

<sup>a</sup>Regimens are described in **Table 1**.
--- indicates not determined.

### Establishment of infection: C3HeB/FeJ mice

The day after aerosol infection, the mean bacterial implantation was 2.03 (SD 0.13) log_10_ CFU/lung (median 2.06 log_10_ CFU/lung) (Figs. 2A, **S9**; **Tables S1**, **S2**). To monitor the infection status during the 6 weeks between infection and the start of treatment, we also determined the lung CFU counts 3 weeks after infection (Table 1). *M. tuberculosis* had multiplied to a mean burden of 6.39 (SD 0.25) log_10_ CFU/lung (median 6.45 log_10_ CFU/lung) (**Table S13**), and unevenly distributed, non-uniform, large lung lesions were just visible (**Fig. S10**). During the subsequent 3 weeks, twenty-two mice died or became moribund and were euthanized prior to treatment initiation. The lungs of these mice were extensively diseased with large, diffuse lesions affecting entire lung lobes (**Fig. S11**), and the mean bacterial burden in these mice was 9.14 (SD 0.21) log_10_ CFU/lung (median 9.11 log_10_ CFU/lung) (**Table S14**). The mice pre-assigned for sacrifice at the Day 0 time point had a mean bacterial burden of 7.46 (SD 0.75) log_10_ CFU/lung (median 7.20, range 6.63-8.78 log_10_ CFU/lung) (**Table S13**), and varying gross lung pathology was also observed (**Fig. S12**). Lung lesions were variable in size and number and non-uniformly distributed. Including the mice that died before the start of treatment, the pre-treatment bacterial burden in the C3HeB/FeJ mice spanned nearly 3 log_10_, ranging from 6.63 to 9.59 log_10_ CFU/lung (Fig. 2A). Among the pre-assigned untreated negative control mice, two died prior to Day 0, and the remaining mice survived 13 to 26 weeks post-infection, with bacterial burdens ranging from 7.74 to 9.29 log_10_ CFU/lung at the time of death (**Table S14**); the lungs of these mice were also extensively diseased (**Fig. S13**). Overall, the course of *M. tuberculosis* infection in this C3HeB/FeJ mouse model was as expected based on previous studies (24, 34, 37, 39).

**Figure 2.**
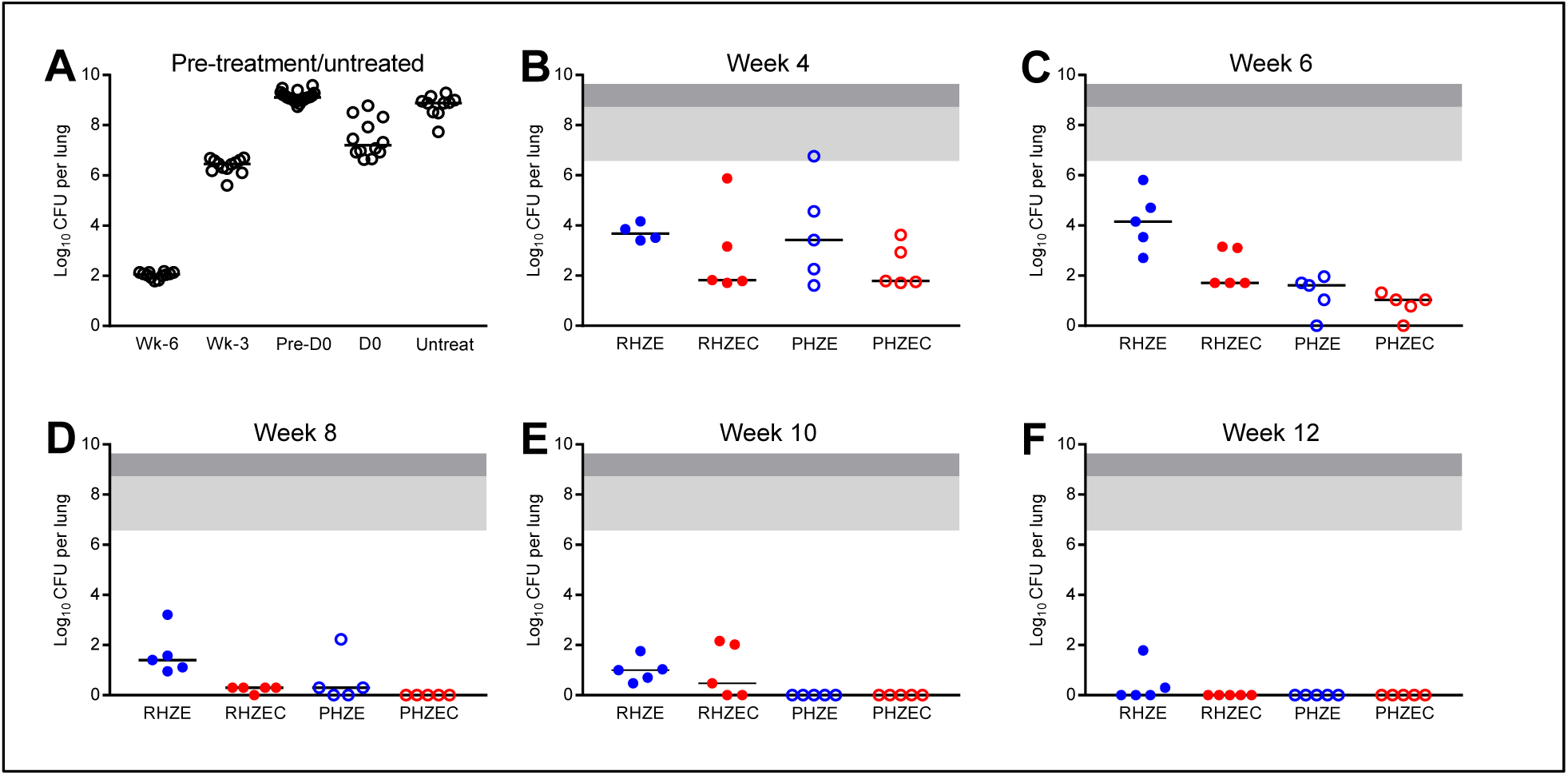
Lung CFU counts of C3HeB/FeJ mice before and during treatment. Lung CFU counts from mice sacrificed the day after infection (“Wk-6”), three weeks after infection (“Wk-3”), and on the day of treatment initiation (“D0”) are shown in **Panel A**, as are the lung CFU counts are for mice that became moribund and were euthanized after Wk-3 but before D0 (“Pre-D0”) and for the untreated negative control mice (“Untreat”). Lung CFU counts after 4, 6, 8, 10, and 12 weeks of treatment are shown in **Panels B**, **C**, **D**, **E**, and **F**, respectively. Treatment regimens are described in Table 1. Due to the variability associated with this model, the lung CFU counts were plotted by individual mouse, with the median value indicated with a black bar. For panels B-F, the area shaded light gray indicates the range of lung CFU counts at the start of treatment in the mice sacrificed at D0, and the area shaded dark gray extends the range to include the lung CFU counts of the mice that died prior to D0. Raw CFU data are presented in **Table S2** (Wk-6), **Table S13** (Wk-3, D0), **Table S14** (Pre-D0, Untreated), and **Tables S15**, **S16**, **S17**, **S18**, and **S19** (Week 4, 6, 8, 10, and 12, respectively).

### Assessment of bactericidal activity: C3HeB/FeJ mice

The RHZE control regimen performed as expected in this model, resulting in killing of 3.5 log_10_ CFU/lung by 4 weeks (Fig. 2B; **Table S15**), with continued bactericidal activity, albeit with more variable CFU counts, over 12 weeks (Fig. 2C-F), resulting in a final mean bacterial burden of 0.42 (SD 0.78) log_10_ CFU/lung (only 2/5 mice were culture-positive at the end of treatment). Gross lung lesions were observed in the RHZE-treated mice up to Week 12 (**Figs. S14-S18**). After four of treatment, there was no statistically significant difference between the bactericidal activity of RHZE and any of the other three regimens (Fig. 2B). As with RHZE, the RHZEC, PHZE, and PHZEC regimens were all associated with bactericidal activity over 12 weeks, but at no point during treatment were statistically significant differences observed in the bacterial burdens of mice receiving these three regimens. Despite the observed variability in bacterial burden, all mice that received PHZEC became culture-negative at Week 6 (Fig. 2C; **Table S16**), and all mice that received PHZE became culture-negative at Week 8 (Fig. 2D; **Table S17**); mice receiving either of these regimens remained culture-negative at Weeks 10 and 12 (Fig. 2D-F; **Tables S18**, **S19**). Mice that received RHZEC became culture-negative at Week 12 (Fig. 2E; **Table S19**). Regimen-associated differences in gross lung pathology became visible after 6 weeks of treatment. As was observed with BALB/c mice, administration of either of the clofazimine-containing regimens was associated with a reduction in visible lesions (**Figs. S14-S18**).

### Assessment of sterilizing activity: C3HeB/FeJ mice

The raw lung CFU data at relapse assessment in mice treated for 6, 8, 10, and 12 weeks are presented in **Tables S20**, **S21**, **S22**, and **S23**, respectively. As expected, approximately half of the mice (11/19) that received RHZE for 12 weeks were culture-positive 6 months after stopping treatment (Table 2; **Fig. S8**). Less than 50% of mice treated with RHZEC, PHZE, or PHZEC experienced culture-positive relapse after treatment durations of 10, 8, and 6 weeks, respectively. The relapse proportion for mice that received RHZE for 12 weeks was significantly higher than the relapse proportion for mice that received ≥8 weeks of PHZEC and ≥10 weeks of treatment with PHZE (p < 0.05). In mice that received PHZEC for 8 and 10 weeks, there was a single relapse, and no relapse occurred at all in mice that received PHZEC for 12 weeks. As was observed with BALB/c mice, the overall rank order of the regimens’ bactericidal activity in C3HeB/FeJ corresponded to their sterilizing activity, *i.e.*, PHZEC > PHZE > RHZEC > RHZE. However, the bacterial burden in the lungs of the culture-positive mice was generally much higher and more variable for the C3HeB/FeJ mice compared to the BALB/c mice (**Fig. S8**). The bacterial burden in the culture-positive C3HeB/FeJ mice ranged from a single CFU to >9 log_10_ CFU/lung.

### Trough serum concentrations of clofazimine and rifapentine

At the sacrifice time points after 4, 8, and 12 weeks of treatment, the trough (about 72 hours post-dose) serum concentrations of clofazimine and rifapentine were measured in mice receiving clofazimine- and rifapentine-containing regimens, respectively (Fig. 3; **Table S24**). For clofazimine, the trough serum concentrations were around 1.5 µg/mL in mice receiving either RHZEC (Fig. 3A) or PHZEC (Fig. 3B), with no statistically significant differences in mean concentrations across time points or between BALB/c and C3HeB/FeJ mice. For rifapentine, differences in trough serum concentrations were observed between mouse strains and between regimens (Figs. 3C-D, **S19**). Rifapentine concentrations were consistently higher in the serum of BALB/c mice compared to C3HeB/FeJ mice. In mice that received PHZE, the mean serum concentration in BALB/c mice was around 8-9 µg/mL at each time point, while the mean serum concentration in C3HeB/FeJ mice was around 4-5 µg/mL (Figs. 3C); the differences were statistically significant at Week 4 (p <0.01) and Week 12 (p <0.05). In mice that received PHZEC (Fig. 3D), mean rifapentine concentrations were higher in BALB/c mice (10.9 µg/mL and 12.1 µg/mL at Weeks 8 and 12, respectively) compared to C3HeB/FeJ mice (7.0 µg/mL and 10.2 µg/mL at Weeks 8 and 12, respectively), though these differences were not statistically significant. In both mouse strains, the mean rifapentine concentrations were higher in mice receiving PHZEC compared those that received PHZE. In C3HeB/FeJ mice that received PHZEC, the mean trough rifapentine concentrations were 30-50% higher than in the C3HeB/FeJ mice that received PHZE; this difference was statistically significant at Week 12 (p <0.01). In BALB/c mice, mean rifapentine concentrations were 25% higher in mice that received PHZEC compared to PHZE, but the difference was not statistically significant.

**Figure 3.**
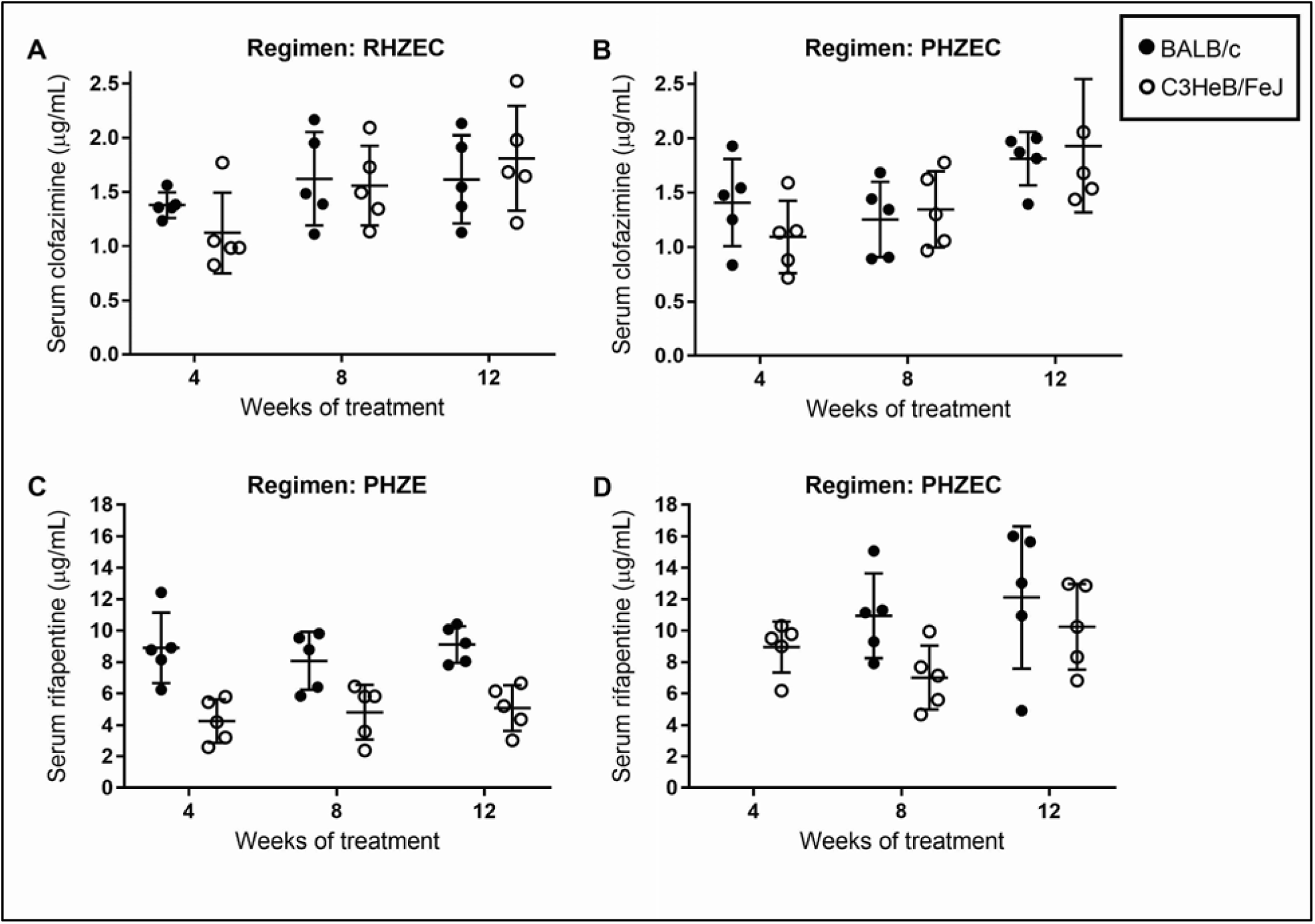
Trough serum clofazimine (Panels A,B) and rifapentine (Panels C,D) concentrations during treatment in BALB/c and C3HeB/FeJ. Serum samples were obtained at sacrifice (about 72 hours after dosing) from mice treated for 4, 8, and 12 weeks. Serum clofazimine levels were determined in mice treated with RHZEC **(Panel A)** and PHZEC (**Panel B**). Serum rifapentine levels were determined in mice treated with PHZE (**Panel C**) and PHZEC (**Panel D**). Individual data points (5 mice per group per time point) are plotted as well as mean and standard deviation values (error bars). For BALB/c mice, the PHZEC Week 4 samples for rifapentine measurement were compromised and discarded. Concentration data for each mouse are presented in **Table S24**.

## DISCUSSION

The main finding of this study is that replacement of rifampin with high-dose rifapentine together with the addition of clofazimine increased the bactericidal and sterilizing activity of the first-line regimen to a significantly greater extent than either modification alone in two pathologically distinct mouse models of TB chemotherapy. While each of these modifications is currently being studied individually in TB patients, our results indicate that superior treatment-shortening effects would be observed if they were combined in the same regimen.

The independent effects of incorporating high-dose rifapentine and clofazimine into the first-line regimen in BALB/c mice were consistent with those observed in previous studies with this model (9, 19). Aggregating the results of relapse assessments from six prior experiments that examined the impact of rifapentine or clofazimine in this model (9, 18, 19, 40) reveals that relapse was observed in 100% (30/30), 40% (23/58), 13% (4/30), and 8% (2/24) of mice treated with RHZ ±E for 12, 16, 20, and 24 weeks, respectively (**Table S25**). The consistency of these results across studies allows estimation of the effect of each modification on the duration of treatment necessary to prevent relapse. In the current study, replacement of rifampin with high-dose rifapentine reduced the percentages of mice relapsing to 30% (6/20) and 6% (1/18) after 8 and 12 weeks of treatment, respectively, indicating that this intervention reduced the treatment needed to prevent a similar number of relapses by at least 8 weeks compared to RHZE. Likewise, the addition of clofazimine reduced the percentages of mice relapsing to 56% (10/18) and 0% (0/20) after 10 and 12 weeks of treatment, respectively, indicating, as prior studies have, that the treatment duration required to prevent the majority of mice from relapsing is 4-6 weeks shorter when clofazimine is added to the first-line regimen at the 12.5 mg/kg dose. Together, these modifications appear to have an additive effect on the anti-TB activity of the first-line regimen. Combining high-dose rifapentine with clofazimine reduced the percentage of mice relapsing to 28% after 6 weeks of treatment, indicating that the combined modifications shortened the treatment needed to obtain similar number of relapses by at least 10 weeks.

Overall, the results in C3HeB/FeJ mice were quite comparable to those observed in BALB/c mice. Significant mouse-to-mouse variability in lung bacterial burden was observed which was due, in large part, to differences in disease progression prior to treatment, as the pretreatment lung bacterial burden ranged from 6.63 to 9.59 log_10_ CFU/lung (Fig. 2A). The magnitude of the variability in bacterial burden within C3HeB/FeJ treatment groups spanned 4 log_10_ CFU/lung in three of the four treatment groups at least once during the first 6 weeks of treatment (Fig. 2B,C), a slightly greater range than what was observed at the start of treatment, consistent with our observations that the lung disease may contribute to progress in the most severely affected mice after initiation of combination chemotherapy. Although there was variability in trough serum concentrations of rifapentine and clofazimine in the C3HeB/FeJ mice (Fig. 3), drug levels did not correlate with lung CFU counts at the individual mouse level (**Fig. S19**, **Tables S15, S17, S24**), and serum rifapentine and clofazimine concentrations were also highly variable in BALB/c mice in which there was much less variability in lung CFU counts (Figs. 1, 3, **S19**). Therefore, in mice that received clofazimine and/or rifapentine, the serum levels of these drugs did not appear to contribute to variability in lung CFU counts during treatment. However, it is possible that tissue-level, and especially lesion-specific, PK differences in C3HeB/FeJ mice contribute to the observed variability in the bactericidal activity associated with each regimen, a concept that has been intimated by numerous studies.

Irwin and colleagues reported that clofazimine monotherapy, administered at 20 mg/kg, had limited bactericidal activity in the lungs of *M. tuberculosis-*infected C3HeB/FeJ mice when treatment was initiated 6 weeks after infection; however, if treatment was initiated 3 weeks after infection, clofazimine had quite potent bactericidal activity (30). The authors linked the diminished activity of clofazimine with the development of hypoxic, caseous necrotic lesions containing extracellular bacteria, suggesting that clofazimine was less active within these lesions. This hypothesis was further supported by studies demonstrating that clofazimine diffuses relatively poorly into caseous necrotic lesions in humans TB (27), does not have bactericidal activity in *ex vivo* caseum from *M. tuberculosis*-infected rabbits (41), and the long-standing observation that clofazimine accumulates in macrophages (26, 27, 42, 43). Rifapentine has been shown to diffuse into necrotic caseous lesions less rapidly than rifampin in a rabbit model of cavitary TB (28), a finding that correlates with clinical data indicating that replacing rifampin with rifapentine added proportionally less activity in patients with cavitary versus non-cavitary TB disease (44). It has also been reported that rifapentine accumulates in macrophages and is relatively more active against intracellular than extracellular *M. tuberculosis* (25). Finally, we previously observed an association between large gross lung lesions in *M. tuberculosis*-infected C3HeB/FeJ mice and reduced rifapentine activity; the bactericidal activity of rifapentine at 10 mg/kg was similar to that of rifampin at 10 mg/kg in mice with the largest gross lesions and highest CFU counts (24). As such gross lung pathology in C3HeB/FeJ mice is associated with the presence of necrotic, caseating granulomatous histopathology (24, 34, 35), these data supported the hypothesis that rifapentine may be less active and/or available in the necrotic, caseating lesions. Therefore, it is possible that the apparent diminished activity of clofazimine and/or rifapentine in the large caseous lesions of some C3HeB/FeJ mice could contribute to the variability in lung CFU counts observed at Week 4, especially among mice receiving rifapentine- or clofazimine-containing regimens. (Fig. 2B), although our data cannot specifically address this issue.

Ultimately, and similar to what was observed in BALB/c mice, both of these drugs did individually add significant bactericidal and sterilizing activity to the first-line regimen in C3HeB/FeJ mice. Possible reasons for this include: the activity of clofazimine and rifapentine was enhanced with co-administration of isoniazid, pyrazinamide, and ethambutol; the anti-TB activity of the other drugs in the regimen allowed the lesions to begin to heal, modifying the necrotic microenvironments and making them more favorable for clofazimine and rifapentine activity; and/or that these drugs reached therapeutic levels at the site of action independent of lung pathology. Interestingly, this overall equivalent activity of the rifapentine-containing regimens was observed between mice strains despite that the trough rifapentine serum levels tended to be lower in the C3HeB/FeJ mice compared to in BALB/c mice (Fig. 3C,D). As the differences were only statistically significant in mice receiving PHZE at the Week 4 and Week 12 time points, it is difficult to interpret the significance, if any, of this difference. Dosing in the C3HeB/FeJ mice was adjusted to account for their increasing body mass over the course of treatment (see Methods and **Table S26**) to ensure that the mice were not under-dosed as they increased in size. Previously, PK differences were not observed in BALB/c and C3HeB/FeJ mice following a single dose of rifapentine alone at 10 mg/kg (9). Thus, further studies are needed to understand any possible long-term PK differences associated with rifapentine between these two strains of mice. Another interesting finding was that in both strains of mice, trough rifapentine levels were higher in mice that received PHZEC compared to PHZE (Fig. 3C,D), suggesting that the presence of clofazimine (which had trough serum levels unaffected by either regimen or mouse strain) may somehow boost rifamycin exposures in the mice, which could in part explain why clofazimine has been shown to contribute significantly more bactericidal activity when added to RHZE than when administered as monotherapy (18, 19, 30, 45–47). Mechanisms of activity aside, this study highlights the essentiality of evaluating the long-term activity of drugs in combination in order to understand the impact of the regimen, as opposed to shorter-term studies and/or studies of single drugs, on treatment outcome.

C3HeB/FeJ mice are more susceptible to *M. tuberculosis* infection than BALB/c mice (22, 24, 33, 34), which was evidenced in this study by the number of mice that became moribund prior to the start of treatment (**Table S14**). This is a limitation of this study in that the C3HeB/FeJ mice with the most severe lung pathology were censored from regimen evaluation. This represents a common quandary with this model. A six-week incubation following aerosol infection is generally necessary to allow for the development of caseous, necrotic lesions in most C3HeB/FeJ mice (22, 24, 33–35). The risk of starting treatment earlier to capture the relatively small proportion of that rapidly succumb to disease is that the majority of the other mice will not develop the pathology that is the hallmark of and rationale for use of this model. In addition to decreasing the bacterial implantation (34), further refinements of this model are needed to limit the heterogeneity in the progression of disease following aerosol infection.

Another limitation of this study is that lung histopathology was not evaluated. We and others have presented histopathological results from this model previously and demonstrated the association between size and extent of caseating lesions and bacterial burden (22, 24, 32, 35). For this study, our primary and secondary endpoints of culture-positive relapse (sterilizing activity) and decline in lung bacterial burden during treatment (bactericidal activity), respectively, relied entirely on the detection of *M. tuberculosis* in the mouse lungs. Therefore, it was most important to homogenize the entire lung for CFU assessment. For C3HeB/FeJ mice, this was particularly important due to the non-uniform and asymmetric development of lung disease in these mice (**Figs. S10-S18**), which confounds estimation of total lung burden if calculated from only a portion of the lung. Accordingly, we could not specifically evaluate the presence and distribution of caseous, necrotic lesions in the lungs at the start of treatment. Since disease in humans is heterogeneous and different pathological states can occur simultaneously in the same host, getting sufficient drug to these different lesions and understanding their activity within such lesions is essential for optimal anti-bacterial activity (23, 32, 48). Therefore, studies in animal models replicating these different pathologies is of interest during preclinical evaluations.

In conclusion, in both BALB/c and C3HeB/FeJ mouse models of TB, replacing rifampin with high-dose rifapentine and adding clofazimine in the first-line regimen resulted in greater bactericidal and sterilizing activity than either modification alone, suggesting that a PHZEC-based regimen may have the potential to significantly shorten the treatment duration for drug-susceptible TB. High-dose rifapentine and clofazimine are each currently being evaluated separately in clinical trials as part of a first-line regimen for TB treatment (ClinicalTrials.gov identifiers NCT02410772 and NCT03474198). The preclinical data presented here provide evidence supporting the clinical evaluation of a regimen combining high-dose rifapentine with clofazimine for treatment of drug-susceptible TB.

## MATERIALS AND METHODS

### Study design

The final scheme of the study is presented in Table 1. The original study protocol included 315 and 328 BALB/c and C3HeB/FeJ mice, respectively. To mitigate losses of C3HeB/FeJ mice, which are more susceptible to *M. tuberculosis* than BALB/c mice (22, 24, 33, 34), we infected an additional 25 mice, for a total of 353 C3HeB/FeJ mice. The primary outcome of this study was the proportion of mice with relapse-free cure, *i.e.,* culture-negative lungs six months after stopping treatment. The secondary endpoint was bactericidal activity, *i.e.*, the decline of *M. tuberculosis* CFU counts during treatment.

### Animals

Female BALB/c mice, aged 5 weeks, were obtained from Charles River Laboratories. Female C3HeB/FeJ mice, aged 4-6 weeks, were obtained from The Jackson Laboratory. All mice were housed in a biosafety level 3 vivarium in individually ventilated cages with sterile wood shavings for bedding. Up to five mice were housed per cage with access to food and water *ad libitum*. Room temperature was maintained at 22-24°C with a 12-hour light/dark cycle. All mice were sacrificed by intentional isoflurane overdose (drop method) followed by cervical dislocation.

### Aerosol infections

*M. tuberculosis* strain H37Rv (American Type Culture Collection strain ATCC 27294) was used for aerosol infections. This stock is susceptible to all drugs used in this study. Bacterial stocks were cultured as previously described (45). To achieve a relatively high-burden infection in BALB/c mice, an actively growing bacterial culture with an optical density at 600 nm (OD_600_) of 1.04 was used directly for aerosol infection. To achieve a low-burden infection in C3HeB/FeJ mice, frozen stock (prepared from a culture with an OD_600_ of 1.04) was thawed and diluted 15-fold in phosphate-buffered saline for aerosol infection. The concentration of each bacterial suspension used for infection was determined as previously described (45) and as detailed in **Table S1**. Mice were infected by aerosol using a full-size Glas-Col Inhalation Exposure System according to the manufacturer’s instructions. BALB/c and C3HeB/FeJ mice were infected in three and four infection runs, respectively. To determine the number of bacteria delivered to the lungs, three mice from each infection run were sacrificed the day after infection, and lung CFU counts were determined as previously described (45) and as detailed in **Table S2**.

### Treatment

BALB/c mice were randomized (stratified by infection run) and assigned to treatment groups and sacrifice cohorts two days before the start of treatment. C3HeB/FeJ mice were randomized (stratified by infection run) nine days before the start of treatment; subsequently, we observed the development of significant variation in body mass of the mice. Therefore, two days before the start of treatment, C3HeB/FeJ mice were randomized again, this time stratified only by body mass (groups of <24, 24 to 25, 25 to <26.5, 26.5 to 28, and >28 g), and then assigned to treatment groups and sacrifice cohorts. Untreated negative control mice were not randomized but followed separately to monitor infection outcome by aerosol infection run. Treatment was initiated on Day 0, which was two and six weeks after infection for BALB/c and C3HeB/FeJ mice, respectively. Three mice from each infection run were sacrificed on Day 0 to determine the lung CFU counts. All pretreatment lung CFU counts were determined as previously described (45).

The regimens evaluated in this study are presented in Table 1. Untreated mice served as the negative control group to verify the virulence of the *M. tuberculosis* infection. The first-line regimen, daily rifampin (R, 10 mg/kg), isoniazid (H, 10 mg/kg), pyrazinamide (Z, 150 mg/kg), and ethambutol (E, 100 mg/kg), was administered as a positive control; these doses for mice are well-established for approximating the area under the plasma concentration-time curve (AUC) produced by recommended doses in humans (49, 50). Three modifications of the standard regimen were evaluated. In regimen RHZEC, clofazimine (C, 12.5 mg/kg) was added to the first-line regimen; although the human pharmacokinetic profile of clofazimine is not well understood, this dose appears to result in steady-state blood concentrations similar to those observed with a 100 mg daily dose in humans (19). In regimen PHZE, rifampin was replaced with high-dose rifapentine (P, 20 mg/kg); this dose in mice approximates the AUC associated with a 1200 mg dose of rifapentine in humans (8, 51). Regimen PHZEC includes both the clofazimine and rifapentine modifications. Treatment was administered for 12 weeks, and all drugs were administered daily (5 days/week, Monday to Friday) by gavage. Rifampin or rifapentine was administered at least one hour before the HZE combination to avoid adverse pharmacokinetic interactions (52, 53). Clofazimine was administered in a third gavage at least 30 minutes after HZE dosing.

Drug formulations were prepared to deliver the drug and dose in a total volume of 0.2 mL per gavage. For the entire duration of treatment, drug formulations for BALB/c mice were prepared based on an average mouse body mass of 20 g. The body mass of C3HeB/FeJ mice continuously increased over time, and the drug concentrations in each formulation were adjusted accordingly, as detailed in **Table S26**. Rifampin, isoniazid, pyrazinamide, and ethambutol were prepared as solutions in distilled water; clofazimine and rifapentine were prepared as a suspension in 0.05% (wt/vol) agarose. For BALB/c mice, the rifapentine suspension was prepared from crushed tablets; for C3HeB/FeJ mice, the rifapentine suspension was prepared from powder for the first 8 weeks of treatment and from crushed tablets for the last 4 weeks of treatment. Rifapentine tablets (Priftin^®^) and powder were provided by Sanofi; all other drugs were purchased from Sigma-Aldrich/Millipore Sigma. Drug stocks were prepared weekly in single or multidrug formulations for administration and were stored at 4°C.

### Assessment of bactericidal activity

Mice were sacrificed 72 hours after the last dose of treatment was administered. Immediately after euthanasia, blood was collected by cardiac puncture. Lungs were dissected from the mice and stored in 2.5 mL phosphate-buffered saline, pH 7.4, at 4°C for at least 48 hours before gross pathology examination. Lungs were homogenized in TenBroeck glass tissue grinders, and undiluted lung homogenate as well as 10-fold serial dilutions of the homogenate were prepared and cultured as described previously (45), with a volume of 0.5 mL per agar plate. The dilution that yielded CFU counts closest to 50 was used to calculate the total CFU/lung, using the CFU counts from either the plain or charcoal-containing plate, whichever had the higher count, for analysis. Plates were incubated at 37°C in sealed plastic bags for at least 4 weeks before the final reading.

### Assessment of sterilizing activity

For mice that received PHZEC, PHZE, RHZEC, and RHZE regimens, relapse assessment began after 6, 8, 10, and 12 weeks of treatment, respectively (Table 1). These treatment durations were selected based on previous data generated in these mouse models with clofazimine or rifapentine with the first-line regimen (8, 9, 19). Mice were sacrificed 6 months after stopping treatment; lungs were removed, examined, and homogenized as described for the assessment of bactericidal activity. One-tenth of the lung homogenate (0.25 mL) was used to prepare four 10-fold serial dilutions. The entire remaining volume of lung homogenate (2.25 mL) was cultured on four agar plates. To allow approximation of lung CFU counts in the lungs of relapsing mice, the 10^-2^ and 10^-4^ dilutions of lung homogenate were also cultured (0.5 mL per plate). Plates were incubated at 37 °C in sealed plastic bags for at least 6 weeks before the final reading. Relapse was defined as having culture-positive lungs, *i.e*., the growth of ≥1 CFU from the lung homogenate.

### Media

*M. tuberculosis* suspensions were grown in 7H9 broth supplemented with a 10% (vol/vol) oleic acid-albumin-dextrose-catalase (OADC) enrichment, 0.5% (vol/vol) glycerol, and 0.1% (vol/vol) Tween 80. Bacterial suspensions (and cognate dilutions) were cultured on 7H11 agar supplemented with 10% (vol/vol) OADC and 0.5% (vol/vol) glycerol (non-selective 7H11 agar). Lung homogenates (and cognate dilutions) were cultured on selective 7H11 agar, *i.e*., 7H11 agar further supplemented with 50 µg/mL carbenicillin, 10 µg/mL polymyxin B, 20 µg/mL trimethoprim, and 50 µg/mL cycloheximide, to selectively cultivate mycobacteria while inhibiting the growth of contaminating bacteria or fungi (54). To detect and limit drug carryover, lung homogenates from at least all treated mice were plated on selective 7H11 agar further supplemented with an adsorbent agent, 0.4% activated charcoal, as described previously (55). BD Difco Middlebrook 7H9 broth powder, BD Difco Mycobacteria 7H11 agar powder, and BD BBL Middlebrook OADC enrichment were obtained from Becton, Dickinson and Company. Remel Microbiology Products Middlebrook 7H11 agar powder, glycerol, and Tween 80 were obtained from Thermo Fisher Scientific, and activated charcoal was obtained from J. T. Baker. All selective drugs were obtained from Sigma-Aldrich/Millipore Sigma. Stock solutions of trimethoprim were prepared in dimethyl sulfoxide, and all other selective drugs were dissolved in distilled water. Drug stocks were filter sterilized, as was the OADC enrichment, prior to use.

### Drug concentration determination

Serum was separate from blood samples as previously described (45). Serum samples were stored at −80°C until analysis. Serum clofazimine and rifapentine concentrations were measured by LC-MS/MS and HPLC, respectively, at the Infectious Disease Pharmacokinetics Laboratory at the University of Florida College of Pharmacy, Gainesville, Florida. The lower limits of quantification for clofazimine and rifapentine were 0.01 and 0.5 µg/mL, respectively.

### Statistical Analyses

All CFU/mL (bacterial suspension) and CFU/lung estimates (*x*) were log-transformed as log_10_(*x* + 1). For samples cultured in parallel on both plain (*i.e.*, charcoal-free) and charcoal-containing selective 7H11 agar, the log_10_ CFU/lung determined from the agar type that yielded the higher CFU/lung estimate was used when calculating mean values and standard deviation (SD). The lower limit of detection was calculated based on the proportion of undiluted lung sample cultured. Comparisons of serum drug concentration data and BALB/c lung CFU data between treatment groups and time points were analyzed using two-way analysis of variance corrected with Tukey’s test for multiple comparisons. Because of the non-Gaussian distribution of the C3HeB/FeJ lung CFU counts during treatment, comparisons of CFU data between treatment groups and time points were analyzed using the non-parametric Kruskal-Wallis test corrected with Dunn’s test for multiple comparisons. The proportions of mice with culture-positive relapse were compared by using Fisher’s exact test. All statistical analyses were performed by using GraphPad Prism 7.02.

## ACKNOWLEDGEMENTS

We thank Jin Lee, Si-Yang Li, Kala Barnes-Boyle, and Jian Xu for help processing the C3HeB/FeJ mice Week 12 relapse samples. Funding for this work was provided by the AIDS Clinical Trials Group, Division of AIDS, National Institutes of Health.

